# Seasonality of protists in the oligotrophic Gulf of Aqaba revealed by imaging flow cytometry

**DOI:** 10.64898/2026.09.09.749164

**Authors:** Herdís G. R. Steinsdóttir, Miguel J. Frada, Derya Akkaynak

## Abstract

Oligotrophic subtropical waters are some of the ocean’s largest continuous ecosystems, yet their protist diversity and community succession remain poorly understood. We examined the protist community in the surface waters of the Gulf of Aqaba at near-daily resolution from March 2024 to July 2025 using an Imaging FlowCytobot. The record revealed a clear seasonal succession. Total cell density increased nearly threefold during winter mixing, driven mainly by an increase in cells smaller than 5 µm. Cells were generally larger during summer, making total biovolume similar in summer and winter. Biovolume reached its highest sustained values during the spring bloom. Despite its low cell density, the summer community was dynamic, with sudden appearances of euglenophytes, flagellates and dinoflagellates at high densities. Coccolithophores and cryptophytes increased during winter, followed by diatoms and ciliates in spring. We also provide a dataset of 31,700 annotated IFCB images.

**Author contributions:** All authors contributed to conceptualization, data collection and writing. HGRS curated the data, implemented the methodology, performed all analyses and visualizations. MJF and DA provided resources, funding, and project administration.

## Introduction

Microbial plankton are abundant and diverse in the ocean, spanning many functional groups, and central among them are the single-celled eukaryotes (protists). These include the phytoplankton that dominate photosynthetic biomass, and heterotrophic microzooplankton that link primary producers to the microbial loop and to higher trophic levels (Calbet and Landry 1999, Worden et al. 2015). Oligotrophic subtropical waters are some of the ocean’s largest continuous ecosystems, contributing an estimated 11–20% of global oceanic net primary production (Dai et al. 2023). Yet protist diversity and community succession in these waters remain poorly understood, a gap of growing consequence these systems are predicted to expand in a warmer future ocean (Polovina et al. 2008, Dai et al. 2023).

To better understand subtropical protist diversity, community succession, and their responses to environmental variability, we examined the surface protist community of the Gulf of Aqaba (GoA) in the northern Red Sea. The GoA is marked by strong seasonal variability in oceanographic conditions. In summer, the water column is stratified and oligotrophic, sharing physical, chemical, and biological characteristics with open-ocean regions (Zarubin et al. 2017, Avrahami et al. 2025b). In winter, deep convective mixing, often reaching 300–700 meters depending on sea-surface cooling, drives a shift to mesotrophic conditions, and, as stratification resets in spring, the availability of nutrients enables the formation of a phytoplankton bloom dominated by diatoms (Zarubin et al. 2017, Avrahami et al. 2025a).

We analyzed the GoA protist community at near-daily resolution over 1.5 years using an Imaging FlowCytobot (IFCB, Olson and Sosik 2007, Sosik and Olson 2007). The IFCB is an automated imaging-in-flow cytometer that combines flow-cytometric particle detection with imaging. Using chlorophyll fluorescence as the trigger for image capture, it records the chlorophyll-containing fraction of the protist community, which includes eukaryotic phytoplankton, mixotrophs, and phagotrophic taxa that carry chlorophyll from ingested prey (hereafter, ‘protists’ refer to this IFCB-detected fraction). We describe the seasonal succession of the protist community and provide a dataset of 31,700 annotated IFCB images. Together, the community description and the curated dataset offer a resource for comparing communities across oligotrophic ecosystems.

## Methods

### Study site and sample collection

Water samples for protist enumeration and imaging were collected approximately daily from 12 March 2024 to 14 July 2025 (291 sampling days), between 10:00 and 12:00 local time, at 2 m depth from the pier of the Interuniversity Institute for Marine Sciences in Eilat (Figure 1 A, station 1; 29°30′06.2″N, 34°55′03.9″E), along the western shore of the GoA. Samples were collected using a weighted 1 L glass container and analyzed live within 15 minutes of collection on the IFCB (McLane Research Laboratories Inc., USA). During a dedicated spring-bloom campaign in 2025, the IFCB was committed to a parallel sampling effort at station A (∼3 km offshore; Figure 1 A) on 18 dates between March and May and was, on those days, unavailable for routine station 1 sampling. These 18 samples from station A (3–7 m depth) were incorporated into the time series (indicated with horizontal line in Figure 2 C). Repeated comparisons in 2017 and 2018 found similar densities at the two stations for three coccolithophore species (supplementary Figure S2), so the two records are treated as one. Environmental data (temperature, nutrients, and chlorophyll *a*) were obtained from the Israel National Monitoring Program in Eilat (https://iui-eilat.huji.ac.il/en/available-data). Chlorophyll *a* was measured daily at station 2, a pier 400 m north of station 1, while temperature and nutrients were measured monthly on a monitoring cruise to station A (Figure. 1 A).

**Figure 1.**
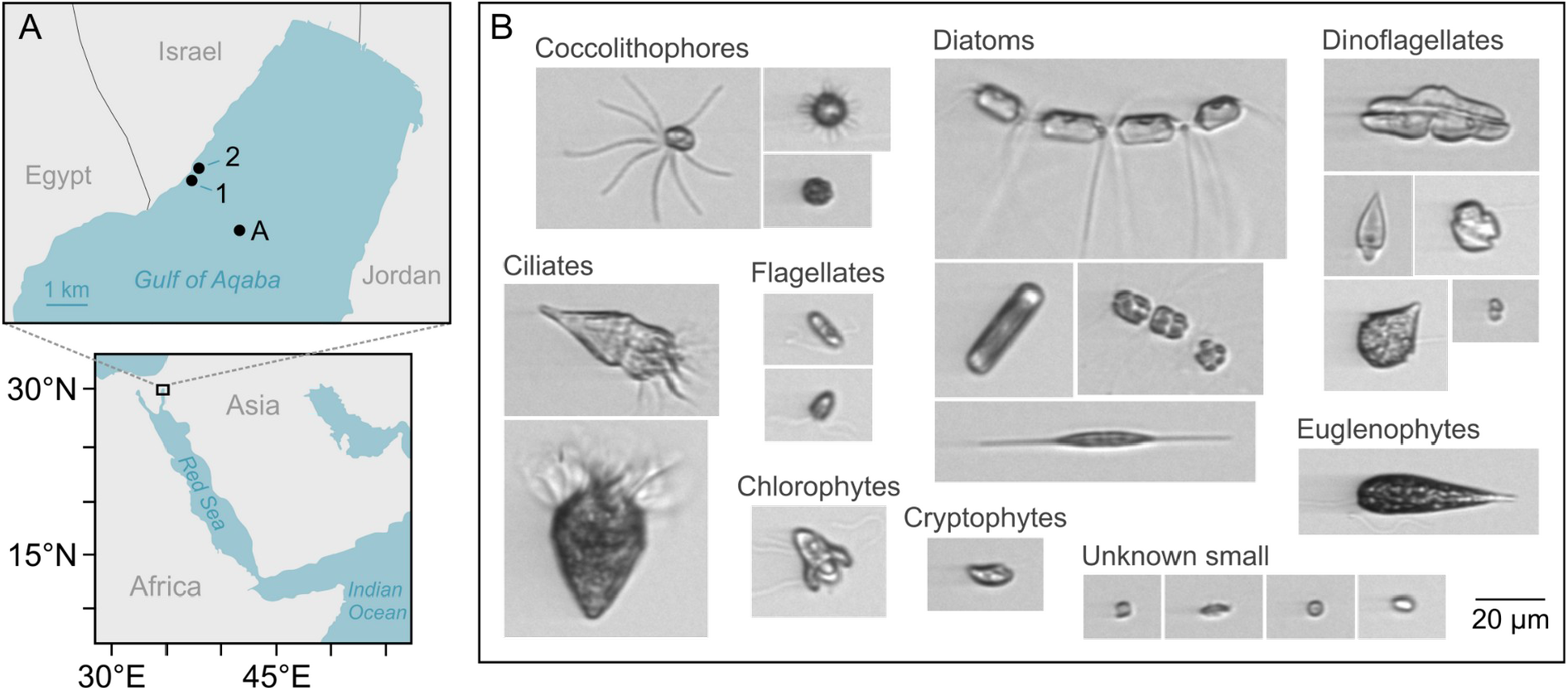
Sampling locations in the Gulf of Aqaba and examples of IFCB image data. (**A**) Map showing the location of routine IFCB sampling at station 1, and the national monitoring program’s measurements of chlorophyll *a* at station 2 (daily) and temperature and nutrients at station A (monthly). (**B**) Example IFCB images grouped by major classes. Scale bar is 20 µm.

**Figure 2.**
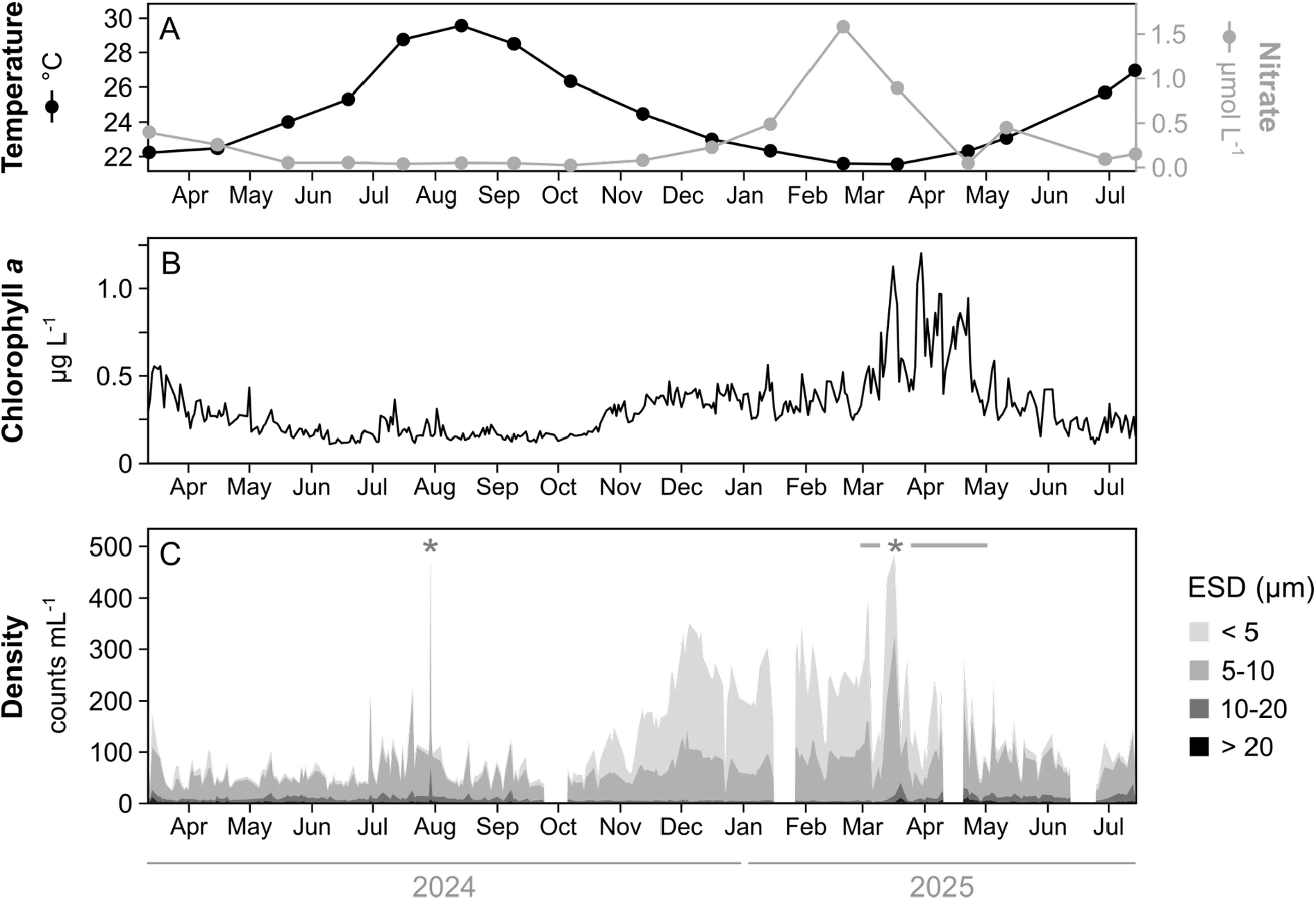
Time series of temperature, nitrate, chlorophyll *a* and protist density in the Gulf of Aqaba. (**A**) Monthly sea-surface temperature (black) and nitrate concentrations (grey). (**B**) Daily chlorophyll *a* concentrations. (**C**) Daily protist density from the IFCB, partitioned by cell size (equivalent spherical diameter, ESD). Asterisks mark two dates where density exceeded the displayed range: 30 July 2024 (841 cells mL^-1^) and 17 March 2025 (644 cells mL^-1^). The horizontal line indicates the period when the time series was supplemented by samples from station A.

### IFCB operation and image acquisition

The IFCB analyses cells and particles between approximately 5–150 μm, and samples were pre-filtered through a 150 μm mesh to prevent clogging. Chlorophyll *a* fluorescence was used as the trigger for image capture, each trigger event generating a grayscale image (region of interest, ROI). A volume of 20 mL was processed per day (four 5 mL runs). The effective volume analyzed, which is lower due to image capturing introducing dead time, ranged between 15.5 and 19.9 mL d^-1^ across the time series and was used to calculate densities and biovolume content. Raw data were written to cloud storage and made available in near-real-time on an online dashboard (www.redsea-ifcb.com). Equivalent spherical diameter (ESD) and biovolume were extracted with the v4 feature pipeline (https://github.com/WHOIGit/ifcb-features, downloaded 15 July 2026). Both were converted from pixels to μm using the instrument’s optical pixel size. Calibration beads of 5.7 µm nominal diameter were imaged for verification occasionally throughout the time series (n = 14 runs) and bead images gave a mean ESD of 5.58 ± 0.16 µm after feature extraction.

### Annotation and training data

We annotated ∼31,700 images from our Gulf of Aqaba time series (archived at Zenodo 10.5281/zenodo.20716677 and browsable at https://ifcbdata.herdis.bio/). Images were annotated using a combination of approaches, including unsupervised clustering of morphologically similar images (MorphoCluster; Schröder et al. 2020), complete annotation of selected samples, targeted searches for under-represented classes, and an iterative loop in which predictions from a preliminary classifier were used to pre-sort images by putative class for manual review. To broaden taxonomic and morphological coverage, we supplemented our annotated data with publicly available IFCB data from other regions, including annotated images from the Mediterranean Sea (Mente et al. 2025), the Northeast Atlantic Ocean (SAMS IFCB Annotated Image Library; https://ifcb-portal.sams.ac.uk/annotatedImages/), and the Californian Ocean Observing System (CalOOS IFCB Dashboard; https://ifcb.caloos.org/dashboard). The compiled dataset consisted of 46,742 annotated images distributed across 96 classes (supplementary Table 1) with at least six images per class. Classes included taxonomic and morphological groups as well as non-living categories such as beads and detritus. For each class, 10% of the images were held out as an evaluation set used for final classifier assessment, and the remaining 90% constituted the development dataset used for iterative model training and testing rounds.

### CNN classifier training and evaluation

A convolutional neural network based on the Inception v3 architecture (Szegedy et al. 2016) was trained to classify the IFCB images. The model was implemented in PyTorch, initialized from ImageNet-pretrained weights, adapted to single-channel grayscale input, and fine-tuned in full with no layers frozen. Input ROIs were padded to 299 × 299 px. During training, images were augmented by random rotations of up to ± 30° and horizontal flipping with a probability of 0.5. The development dataset was divided by stratified random sampling into 80% for training and 20% for validation, preserving the relative representation of each class. The model was optimised using AdamW with a learning rate of 0.0001 and weight decay of 0.0001, minimizing cross-entropy loss. The auxiliary classifier of Inception v3 contributed to the loss with a weight of 0.4. The learning rate was reduced on plateau by a factor of 0.1 after four epochs without improvement in validation loss. Training ran for 150 epochs with a batch size of 128 on an NVIDIA GeForce RTX 5070 Ti GPU. The epoch with the highest weighted F1 on the validation set (epoch 147, accuracy 97.8%) was retained. Class-specific confidence thresholds were selected from precision-recall curves calculated on the validation set and performance was assessed on the held-out evaluation set and is reported as per-class precision and recall (supplementary Table S1, supplementary Figure S1). Predictions below thresholds were assigned as unclassified.

### Scanning Electron Microscopy

Seawater samples (2–4 L) were pre-screened through a 200 μm mesh and gently filtered using a vacuum pump system (<150 mmHg) onto polycarbonate membranes (Whatman; 0.8 μm pore size, 47 mm diameter). The membranes were mounted on aluminum stubs, sputter-coated with gold–palladium (∼20 nm) and analyzed using a scanning electron microscope (Phenom Pro benchtop and JEOL 6700). Cells were identified and enumerated as described in Keuter et al. (2023), and densities were calculated from the membrane area examined and the volume filtered. The coccolithophores *Umbilicosphaera sibogae* and *Gephyrocapsa huxleyi* were counted for comparison with IFCB estimates obtained from aliquots of the same sample.

### Community analysis

For multivariate community analysis on IFCB classes, non-living, non-protist and low confidence classes were excluded, including aggregates, beads, bubbles, debris, blur, cut-off images, fecal pellets, copepod nauplii, unknown small, and unclassified objects. ‘Unknown small’ and ‘Unclassified’ were retained in visualisations of density and biovolume over time. Daily observations were aggregated into a day-by-class abundance matrix of cells mL^-1^. Days containing fewer than 100 ROIs in total were excluded, as were classes detected on fewer than 5% of the days. Counts were square-root transformed before clustering and ordination to reduce the influence of the most abundant classes. Bray-Curtis dissimilarities were calculated between days, which were then grouped using hierarchical clustering with average linkage (UPGMA). Six clusters were selected based on mean silhouette width evaluated for k 2–10. Community structure was visualized using two-dimensional non-metric multidimensional scaling (NMDS) based on the same dissimilarity matrix. Classes contributing most strongly to differences among clusters were identified using SIMPER. Turnover was calculated as the Bray-Curtis dissimilarity between successive observations separated by no more than three days using untransformed abundances to retain changes in both total abundance and composition. Temporal variation was summarized using a 14-day tolling median, and gaps longer than six days were shown as breaks in the time series. All analyses were conducted at full class resolution and classes were aggregated into broader taxonomic groups for visualization only (Figure 4 and 5). Analysis was performed in R v4.3.3.

## Results and Discussion

The IFCB time series ran from 12 March 2024 until 14 July 2025 and recorded a complete stratification-mixing cycle. The warm, oligotrophic period from June to October was characterized by low concentrations of nitrate (< 0.2 µmol L-1) and chlorophyll *a* (< 0.5 µg L-1; Figure 2 A,B). During the autumn months, cooling of the surface waters eroded the thermocline and caused the water column to mix, bringing nutrients to the surface. The concentrations of chlorophyll *a* rose as the water column mixed, then peaked during a spring phytoplankton bloom in March as stratification returned. In 2024, the concentrations of chlorophyll *a* peaked at 0.6 µg L-1, whereas the bloom was more pronounced in 2025 when concentrations reached 1.2 µg L-1 (Figure 2 A,B).

The density of protists followed the same broad seasonal pattern as chlorophyll (Figure 2). During the stratified period from June to October, densities averaged 75 ± 88 cells mL-1 (mean ± SD; n=94 days) and increased to 221 ± 75 cells mL-1 during the mixing period from November to February. The large variation during summer was primarily due to a single day in July when density reached 841 cell mL-1 (Figure 2 C).

### Cell size and IFCB detection

The increase in protists density during winter was driven by the smallest cells. Cells with an equivalent spherical diameter (ESD) below 5 µm averaged 13 ± 14 cells mL-1 in summer and increased to 144 ± 53 cells mL-1 in winter (Figure 2 C). Large cells were rare throughout the time series and did not follow the same seasonal pattern. Cells with an ESD above 10 µm averaged 8 ± 8 cells mL-1 during summer and decreased to 3.4 ± 0.9 cells mL-1 during winter. Their density never exceeded 72 cells mL-1 and remained below 6.3 cells mL-1 between November and February. Across the time series, between 68 and 99% of imaged cells had an ESD below 10 µm, near the size at which morphological features become difficult to resolve in IFCB images (Figure 1 B). The cells that could not be confidently assigned to a taxonomic class were placed in the catch-all class ‘Unknown small’, which accounted for 13 to 80% of cells across samples (Figure 4 A).

To assess detection, we compared IFCB counts with scanning electron microscopy for two coccolithophore species of differing size and morphology (Figure 3). The two methods resulted in similar densities for the larger *Umbilicosphaera sibogae* (10–20 µm, n = 8), whereas the IFCB recorded approximately seven times fewer cells of the smaller *Gephyrocapsa huxleyi* (3–7 µm; n = 35; median sample-level ratio). A similar 20-fold-discrepancy has been reported between IFCB and microscopy counts of single cells of the chrysophyte *Uroglena* (∼10 µm; Gifford et al. 2024).

**Figure 3.**
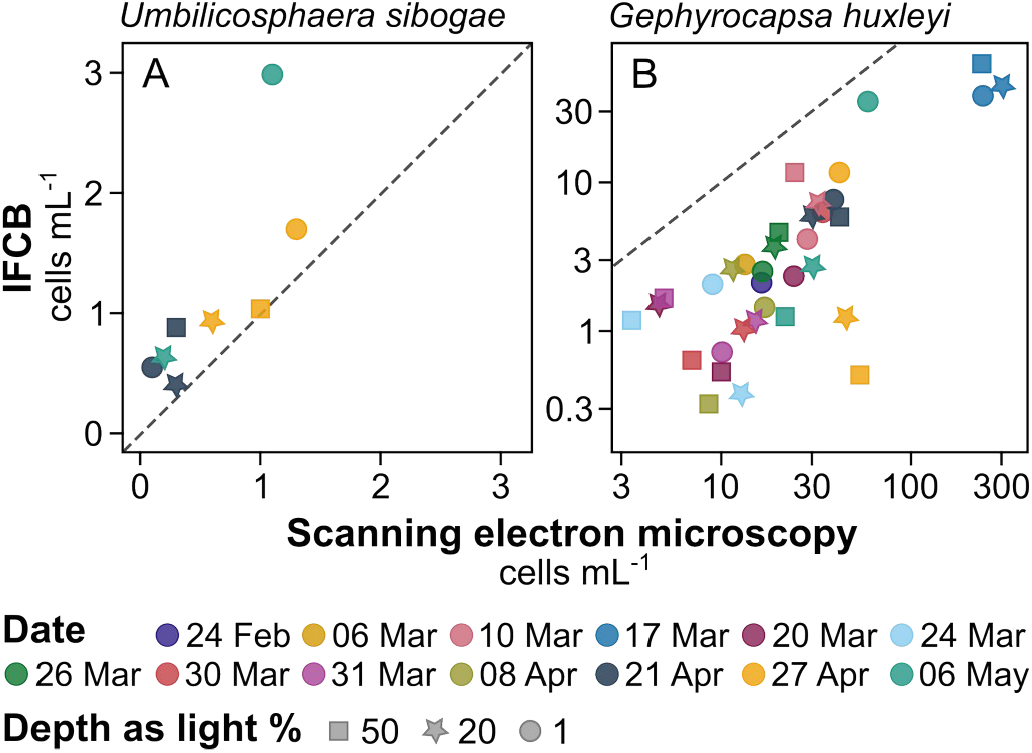
Comparison of IFCB and scanning electron microscopy cell counts for two coccolithophore species of differing size and morphology: (**A**) *Umbilicosphaera sibogae*, (**B**) *Gephyrocapsa huxleyi*. Samples were collected during the bloom in 2025 and each point shows a sample, colored by date. The shape indicates sampling depth as light %. The dashed line is the 1:1 relationship. Note the logarithmic axes in (B).

How well the IFCB detects small cells may also vary with season. In the GoA, daily integrated PAR at 8-m depth was about 45% higher in summer than in winter (Steinsdóttir et al. 2026), which may have reduced cellular chlorophyll content (Stambler 2006) and thereby weakened fluorescence. Since we did not quantify this effect, we cannot determine how much of the summer minimum in small cells was due to lower cell densities and how much was due to reduced detection efficiency. The observed increase in small cells during winter is, however, consistent with earlier work in the GoA (Lindell and Post 1995), and recent molecular data show a winter community dominated by small chlorophytes (Avrahami et al. 2025b). Using light scattering as the trigger for IFCB imaging did not improve detection since it recorded many inorganic particles, potentially pier-associated resuspended material, at the cost of chlorophyll-containing cells.

Despite the challenges posed by the relatively few large cells, the IFCB still resolves many protists to class and even genus level and complements previous work in the GoA nicely. It records cell size and morphology, and reveals seasonal succession and short-lived pulses that less-frequent sampling would have missed. Trends among the major groups are shown in Figure 4 and 5, and all class-level time series are available at https://ifcb-pier-herdissteins.pythonanywhere.com.

**Figure 4.**
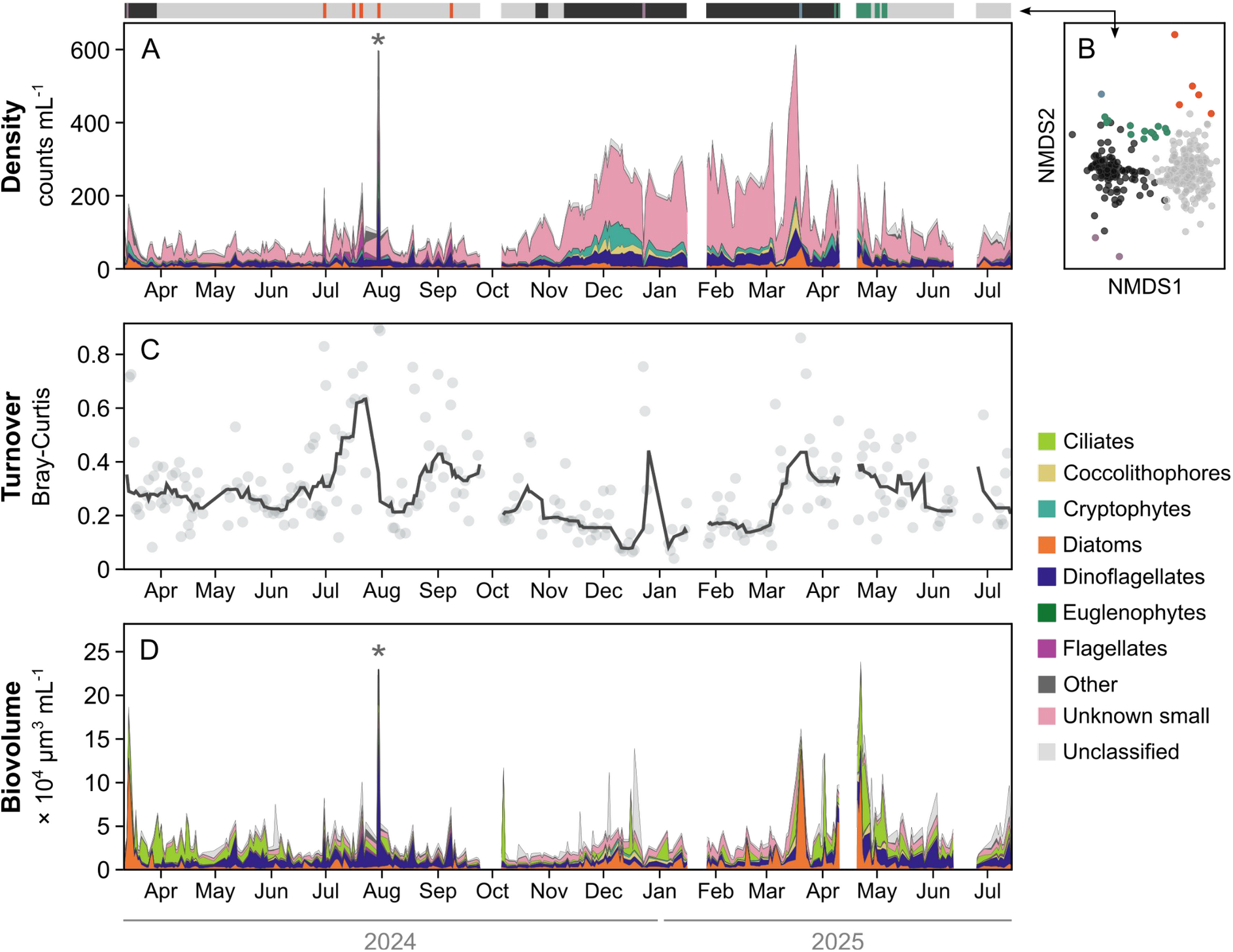
Seasonal dynamics of the protist community in the GoA. (A) Daily density stacked by class. The bar above the panel shows each day’s assignment to the community clusters in (B), colored by cluster. (B) NMDS ordination of samples based on Bray-Curtis dissimilarity, excluding ‘Unknown small’. (C) Daily community turnover between consecutive samples (Bray-Curtis dissimilarity). The line indicates a 14-day rolling median. (D) Daily estimated biovolume stacked by class. Asterisks in (A) and (D) mark 30 July 2024, where values exceed the displayed range (Density: 841 cells mL ^-1^, Biovolume: 32 × 10^4^ µm^3^ mL^-1^).

**Figure 5.**
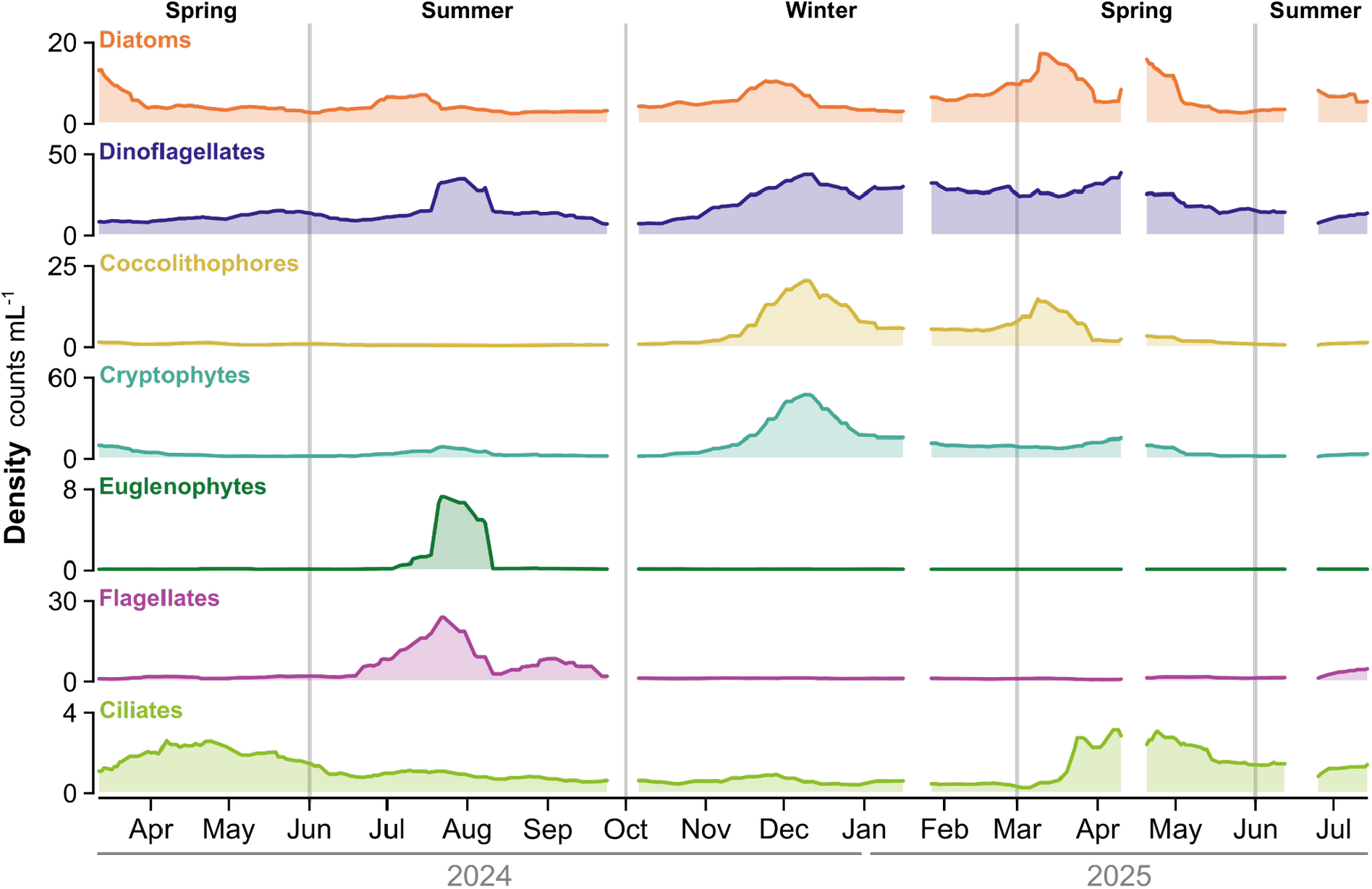
Smoothed density of major protist groups in the GoA. Lines are centered 21-day rolling means calculated from observations within 10 days before and after each date (minimum three observations). Note variation in y-axis between groups.

### Seasonal succession of protists in the GoA

Most protist classes demonstrated seasonality and the summer and winter assemblages formed separate clusters (Figure 4 A,B). A smoothed group-level view of the time series illustrates the succession of the major groups (Figure 5). Coccolithophores and cryptophytes were scarce during summer but increased as mixing began in November, consistent with previous microscopy observations (Al-Najjar et al. 2007; Keuter et al. 2023). They averaged 8 and 19 cells mL-1 during winter, compared with 0.5 and 3 cells mL-1 during summer, respectively. In contrast, flagellates (a morphological group) and euglenophytes were largely confined to the summer, whereas dinoflagellates were abundant throughout the year, their density increasing from an average of 13 cells mL-1 in summer to 27 cells mL-1 in winter. The presence of flagellates, euglenophytes and dinoflagellates (classes in which mixotrophy is common) during summer is consistent with earlier molecular work, which suggests that phago-mixotrophic groups are an important part of the summer community in the GoA (Avrahami et al. 2025b). Diatoms were most abundant in the spring, averaging 8 and 12 cells mL-1 in March of 2024 and 2025, respectively. Ciliate density increased in the weeks after the diatom peak in both years, consistent with a grazer’s response to the bloom. In 2025, several samples from the spring bloom formed a separate cluster (Figure 4 B).

Biovolume revealed a slightly different seasonal pattern from cell density. Total biovolume was generally highest during March and April, when it ranged 13,100 to 237,100 µm3 mL-1 (Figure 4 D). Although diatoms and ciliates accounted for only 9% of cells during these months, they contributed 59% of the biovolume. And despite the nearly threefold difference in cell density between summer and winter, the average biovolume was similar between the two seasons (29,700 and 31,300 µm3 mL-1, respectively). This stability was driven primarily by dynamics within the dinoflagellates. Their density was twice as high in winter, whereas their biovolume was more than twice as high in summer, contributing 32% of the total biovolume. Thus, despite low cell densities, the summer community contained, on average, larger cells and had a total biovolume comparable to that in winter.

Community composition also changed within seasons. Turnover between successive samples was generally higher in spring and summer than in winter (Figure 4 C). Five samples from late summer formed a separate cluster, driven mainly by sudden appearances of euglenophytes and several flagellate and dinoflagellate classes. The strongest pulse occurred on 30 July 2024, when total density reached 841 cells mL-1, higher than in any spring-bloom sample. Samples collected before and after it returned to the usual summer assemblage. The cause of these pulses cannot be resolved from these observations alone. However, a recent modeling effort suggests that horizontal transport from the southern, more-productive part of the GoA contributes to the spring bloom in the northern GoA (Berman and Gildor, 2022). A similar advective process might be at play in late summer and contribute to the rapid community changes observed then. Whatever the cause, the oligotrophic summer community was neither uniform nor static, and substantial changes occurred over the timescale of a few days.

The IFCB record reveals pronounced seasonal succession in the protist assemblage of the GoA. Summer flagellates and dinoflagellates gave way to cryptophytes and coccolithophores during winter, followed by diatoms and ciliates in spring. Cell density changed markedly across seasons, being highest in winter and spring. In contrast, total biovolume was comparatively stable over time, reflecting the abundance of small cells in winter and the presence of fewer, larger cells in summer. Turnover was generally higher in spring and summer than in winter, with several short-lived pulses of increased densities recorded in the late summer. Finally, the accompanying dataset of 31,700 annotated images provides a resource for improving IFCB classifications in oligotrophic waters.

## Acknowledgements

We are grateful to Gil Koplovitz for assistance with sampling and scanning electron microscopy, and Liraz Levy for sampling and logistical support. help with field sampling, analysis, and logistical support. We thank Yonni Shaked, Amatzia Genin, and the Israel National Monitoring Program in Eilat for providing environmental data. We also thank Yoav Avrahami for help with image annotation and Grigory Solomatov for discussions on classifier optimization.

## Data availability

The annotated image dataset underlying the classifier, comprising 31,700 images assigned to 96 classes, is archived at Zenodo (https://doi.org/10.5281/zenodo.20716677) and can be browsed at https://ifcbdata.herdis.bio.

## Notes

https://ifcbdata.herdis.bio

https://ifcb-pier-herdissteins.pythonanywhere.com

### Competing Interest Statement

The authors have declared no competing interest.

