## Supplementary material for "Seasonality of protists in the oligotrophic Gulf of Aqaba revealed by imaging flow cytometry"

**Table 1.** Per-class performance of the 96-class classifier on the held-out evaluation set. *n*, total number of annotated images; Thr, class-specific probability threshold; *P*, precision; *R*, recall; $F_1$, F-score (harmonic mean of *P* and *R*). Classes shaded in orange contain fewer than 100 annotated images and those shaded in blue have $F_1$ below 0.90. Results for the rare classes should be interpreted with caution.

| Class | *n* | Thr. | Prec. | Rec. | F1 |
|---|---|---|---|---|---|
| aggregate | 803 | 0.50 | 0.91 | 0.90 | 0.905 |
| beads | 664 | 0.99 | 1.00 | 1.00 | 1.000 |
| beads_multiple | 121 | 0.66 | 1.00 | 1.00 | 1.000 |
| blur_faint | 1,112 | 0.97 | 0.96 | 0.96 | 0.960 |
| bubble | 61 | 0.99 | 0.92 | 1.00 | 0.958 |
| centrohelea | 6 | 0.95 | 0.50 | 1.00 | 0.667 |
| chlorophyta_pyramimonas | 574 | 0.83 | 0.99 | 0.99 | 0.990 |
| chlorophyta_pyramimonas_small | 597 | 0.95 | 1.00 | 1.00 | 1.000 |
| chrysophyceae_dinobryon | 2,336 | 0.94 | 1.00 | 1.00 | 1.000 |
| ciliate_mesodinium | 566 | 0.50 | 1.00 | 0.99 | 0.995 |
| ciliate_mix | 1,777 | 0.39 | 0.98 | 0.99 | 0.985 |
| ciliate_tiarina | 43 | 0.99 | 1.00 | 1.00 | 1.000 |
| ciliate_tintinnid | 244 | 0.52 | 0.92 | 0.94 | 0.930 |
| cocco_01 | 191 | 0.70 | 1.00 | 1.00 | 1.000 |
| cocco_calciopappus_michaelsarsia | 125 | 0.93 | 0.96 | 1.00 | 0.980 |
| cocco_calciosolenia | 247 | 0.99 | 1.00 | 1.00 | 1.000 |
| cocco_gephyrocapsa_huxleyi | 826 | 0.24 | 1.00 | 0.98 | 0.990 |
| cocco_ophiaster | 164 | 0.98 | 1.00 | 1.00 | 1.000 |
| cocco_rhabdosphaera | 278 | 0.98 | 1.00 | 1.00 | 1.000 |
| cocco_syracosphaera | 110 | 0.83 | 1.00 | 0.95 | 0.974 |
| cocco_syracosphaera_pulchra | 178 | 0.99 | 1.00 | 1.00 | 1.000 |
| cocco_umbilicosphaera_sibogae | 146 | 0.70 | 1.00 | 1.00 | 1.000 |
| copepod_nauplii | 105 | 0.93 | 1.00 | 1.00 | 1.000 |
| cryptophyte | 3,604 | 0.83 | 0.99 | 0.99 | 0.990 |
| cutoff_half | 143 | 0.89 | 1.00 | 0.69 | 0.817 |
| debris_inorg | 316 | 0.74 | 0.93 | 0.79 | 0.854 |
| diatom_01 | 33 | 0.87 | 0.83 | 0.71 | 0.765 |
| diatom_02 | 44 | 0.99 | 1.00 | 1.00 | 1.000 |
| diatom_03 | 52 | 0.82 | 1.00 | 1.00 | 1.000 |
| diatom_bacteriastrum_furcatum | 334 | 0.96 | 1.00 | 0.80 | 0.889 |
| diatom_bacteriastrum_jadranum | 635 | 0.92 | 0.98 | 0.97 | 0.975 |
| diatom_bacteriastrum_other | 86 | 0.96 | 1.00 | 0.76 | 0.864 |
| diatom_centric_medium_large | 171 | 0.91 | 1.00 | 1.00 | 1.000 |
| diatom_centric_small | 458 | 0.50 | 1.00 | 0.97 | 0.985 |
| diatom_cerataulina | 40 | 0.83 | 1.00 | 0.88 | 0.936 |
| diatom_chaetoceros | 1,370 | 0.81 | 0.98 | 0.96 | 0.970 |
| diatom_chaetoceros_cyst | 190 | 0.71 | 0.97 | 0.92 | 0.944 |
| diatom_corethron | 92 | 0.99 | 1.00 | 1.00 | 1.000 |
| diatom_cylindrotheca | 1,275 | 0.90 | 0.99 | 0.98 | 0.985 |
| diatom_eucampia | 61 | 0.14 | 0.92 | 0.92 | 0.920 |
| diatom_guinardia | 451 | 0.91 | 1.00 | 1.00 | 1.000 |
| diatom_helicotheca | 8 | 0.99 | 1.00 | 0.50 | 0.667 |
| diatom_hemiaulus | 54 | 0.47 | 1.00 | 0.91 | 0.953 |
| diatom_leptocylindrus | 669 | 0.57 | 0.99 | 0.96 | 0.975 |
| diatom_licmophora | 246 | 0.94 | 1.00 | 0.94 | 0.969 |
| diatom_nanoneis | 98 | 0.98 | 1.00 | 1.00 | 1.000 |
| diatom_pennate_long | 41 | 0.93 | 1.00 | 0.75 | 0.857 |
| diatom_pennate_mix | 1,800 | 0.74 | 1.00 | 0.98 | 0.990 |
| diatom_pennate_thin_uncertain | 260 | 0.98 | 0.95 | 0.75 | 0.838 |
| diatom_pleurosigma | 175 | 0.08 | 1.00 | 0.97 | 0.980 |
| diatom_pseudonitzschia | 682 | 0.83 | 0.97 | 0.90 | 0.934 |
| diatom_rhizosolenia_proboscia | 272 | 0.70 | 0.95 | 0.98 | 0.965 |
| diatom_skeletonema | 398 | 0.93 | 1.00 | 0.94 | 0.969 |
| diatom_striatella | 305 | 0.70 | 1.00 | 1.00 | 1.000 |
| diatom_thalassionema | 223 | 0.99 | 1.00 | 1.00 | 1.000 |
| diatom_tropidoneis_plagiotropis_maybe | 46 | 0.96 | 1.00 | 0.78 | 0.876 |
| dictyochales_silicoflagellates | 810 | 0.99 | 0.99 | 1.00 | 0.995 |
| dino_akashiwo | 107 | 0.99 | 0.95 | 1.00 | 0.974 |
| dino_balechina | 8 | 0.99 | 1.00 | 1.00 | 1.000 |
| dino_corythodinium | 33 | 0.14 | 1.00 | 1.00 | 1.000 |
| dino_dinophysis | 372 | 0.99 | 1.00 | 0.97 | 0.985 |
| dino_gonyaulax | 38 | 0.99 | 1.00 | 0.50 | 0.667 |
| dino_gonyaulax_long | 7 | 0.35 | 1.00 | 1.00 | 1.000 |
| dino_gyrodinium | 184 | 0.92 | 1.00 | 0.97 | 0.985 |
| dino_kapelodinium | 191 | 0.99 | 1.00 | 0.87 | 0.930 |
| dino_karenia_papilionacea | 189 | 0.89 | 1.00 | 0.95 | 0.974 |
| dino_karenia_round | 21 | 0.66 | 1.00 | 0.25 | 0.400 |
| dino_large_dark | 21 | 0.99 | 1.00 | 1.00 | 1.000 |
| dino_mix | 2,450 | 0.98 | 0.99 | 0.95 | 0.970 |
| dino_oxytoxum_large | 73 | 0.99 | 1.00 | 1.00 | 1.000 |
| dino_oxytoxum_small | 130 | 0.95 | 1.00 | 0.96 | 0.980 |
| dino_prorocentrum | 370 | 0.70 | 0.99 | 0.99 | 0.990 |
| dino_protoperidinium | 167 | 0.99 | 1.00 | 0.88 | 0.936 |
| dino_scrippsiella | 712 | 0.83 | 1.00 | 0.99 | 0.995 |
| dino_small_long | 842 | 0.56 | 0.99 | 0.98 | 0.985 |
| dino_small_squiggle | 1,146 | 0.57 | 0.99 | 0.96 | 0.975 |
| dino_torodinium_katodinium | 488 | 0.94 | 1.00 | 0.99 | 0.995 |
| dino_tripos | 4,963 | 0.83 | 1.00 | 1.00 | 1.000 |
| dino_warnowia_cochlodinium | 358 | 0.14 | 1.00 | 1.00 | 1.000 |
| euglenophytes | 625 | 0.83 | 1.00 | 1.00 | 1.000 |
| fecalpellet | 324 | 0.98 | 1.00 | 0.94 | 0.969 |
| flagellate_01 | 119 | 0.03 | 1.00 | 1.00 | 1.000 |
| flagellate_02 | 897 | 0.94 | 1.00 | 1.00 | 1.000 |
| flagellate_03 | 192 | 0.90 | 1.00 | 0.90 | 0.947 |
| flagellate_04 | 255 | 0.96 | 0.98 | 0.98 | 0.980 |
| haptophyta_chrysochromulina_parkeae | 496 | 0.93 | 0.99 | 1.00 | 0.995 |
| haptophyta_chrysochromulina_putative | 248 | 0.95 | 0.96 | 0.84 | 0.896 |
| haptophyta_phaeocystis_colonies | 383 | 0.99 | 1.00 | 0.94 | 0.969 |
| radiolaria | 200 | 0.14 | 0.98 | 1.00 | 0.990 |
| radiolaria_spherical | 13 | 0.99 | 1.00 | 1.00 | 1.000 |
| two_tintinnid_chaetoceros | 7 | 0.99 | 1.00 | 1.00 | 1.000 |
| unknown_01 | 9 | 0.99 | 1.00 | 1.00 | 1.000 |
| unknown_02 | 100 | 0.99 | 1.00 | 0.95 | 0.974 |
| unknown_03 | 224 | 0.97 | 1.00 | 0.98 | 0.990 |
| unknown_04 | 61 | 0.06 | 1.00 | 1.00 | 1.000 |
| unknown_small | 3,003 | 0.75 | 0.99 | 0.97 | 0.980 |

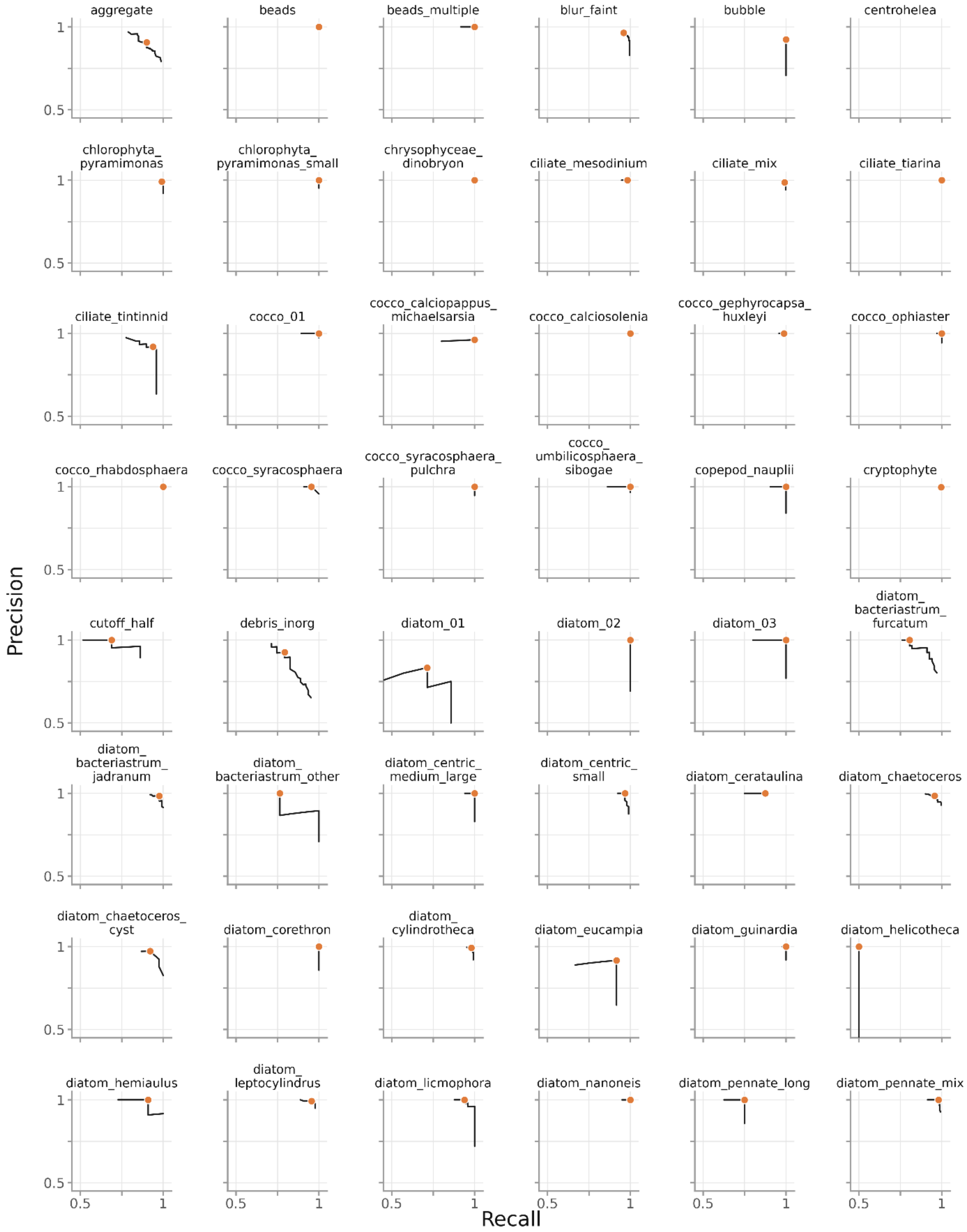
aggregate
beads
beads_multiple
blur_faint
bubble
centrohelea
chlorophyta_ pyramimonas
chlorophyta_ pyramimonas_small
chrysophyceae_ dinobryon
ciliate_mesodinium
ciliate_mix
ciliate_tiarina
ciliate_tintinnid
cocco_01
cocco_calciopappus_ michaelsarsia
cocco_calciosolenia
cocco_gephyrocapsa_ huxleyi
cocco_ophiaster
cocco_rhabdosphaera
cocco_syracosphaera
cocco_syracosphaera_ pulchra
cocco_ umbilicosphaera_ sibogae
copepod_nauplii
cryptophyte
cutoff_half
debris_inorg
diatom_01
diatom_02
diatom_03
diatom_ bacteriastrum_ furcatum
diatom_ bacteriastrum_ jadranum
diatom_ bacteriastrum_other
diatom_centric_ medium_large
diatom_centric_ small
diatom_cerataulina
diatom_chaetoceros
diatom_chaetoceros_ cyst
diatom_corethron
diatom_ cylindrotheca
diatom_eucampia
diatom_guinardia
diatom_helicotheca
diatom_hemiaulus
diatom_ leptocylindrus
diatom_licmophora
diatom_nanoneis
diatom_pennate_long
diatom_pennate_mix
Precision
Recall
0.5
1

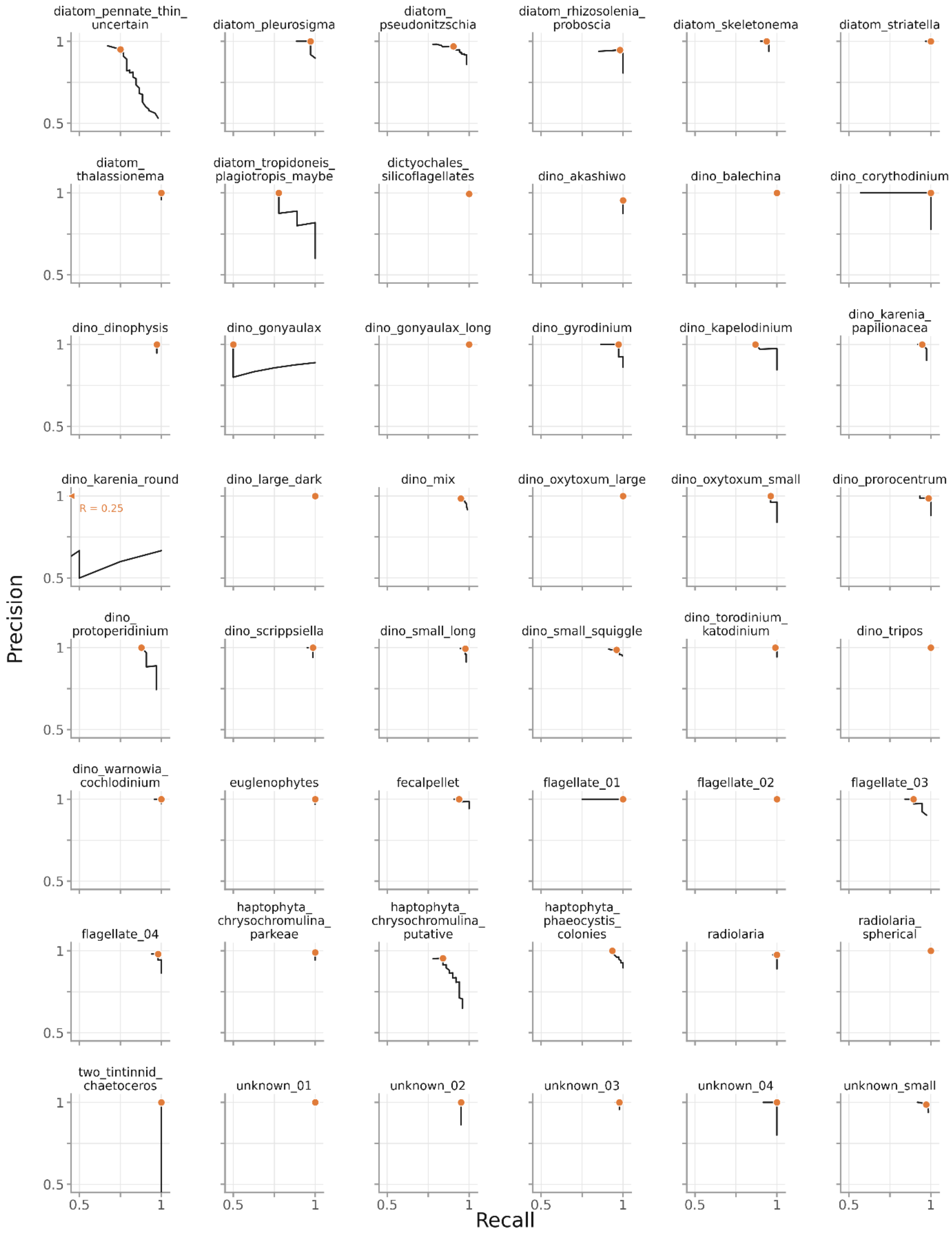


**Figure S1.** Class-specific precision-recall curves. Orange point shows the performance at the selected threshold.

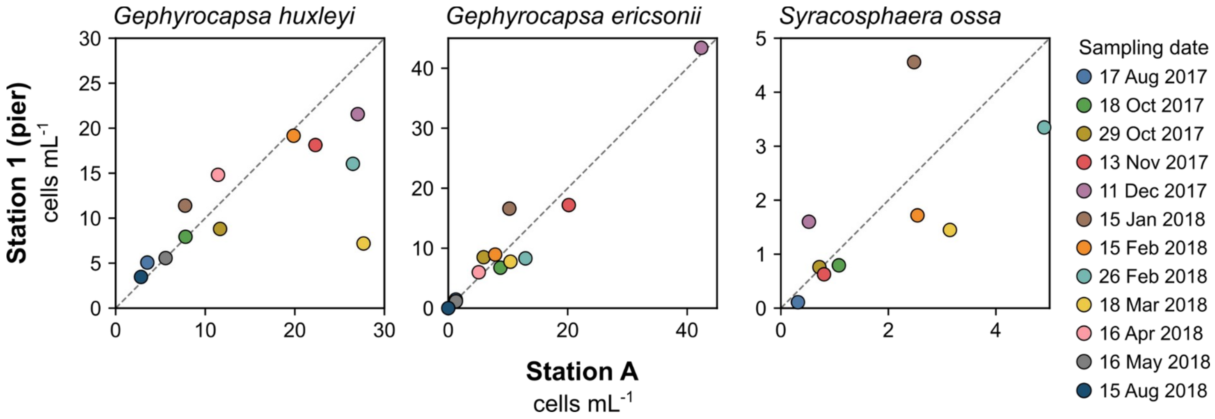


**Figure S2.** Comparison of cell densities at station 1 (pier) and station A, about 3 km offshore, for three coccolithophore species: *Gephyrocapsa huxleyi*, *Gephyrocapsa ericsonii* and *Syracosphaera ossa*. Samples at both stations were collected from surface waters within one hour of each other on 12 dates between August 2017 and August 2018. Cells were counted by scanning electron microscopy. The dashed line is the 1:1 relationship.

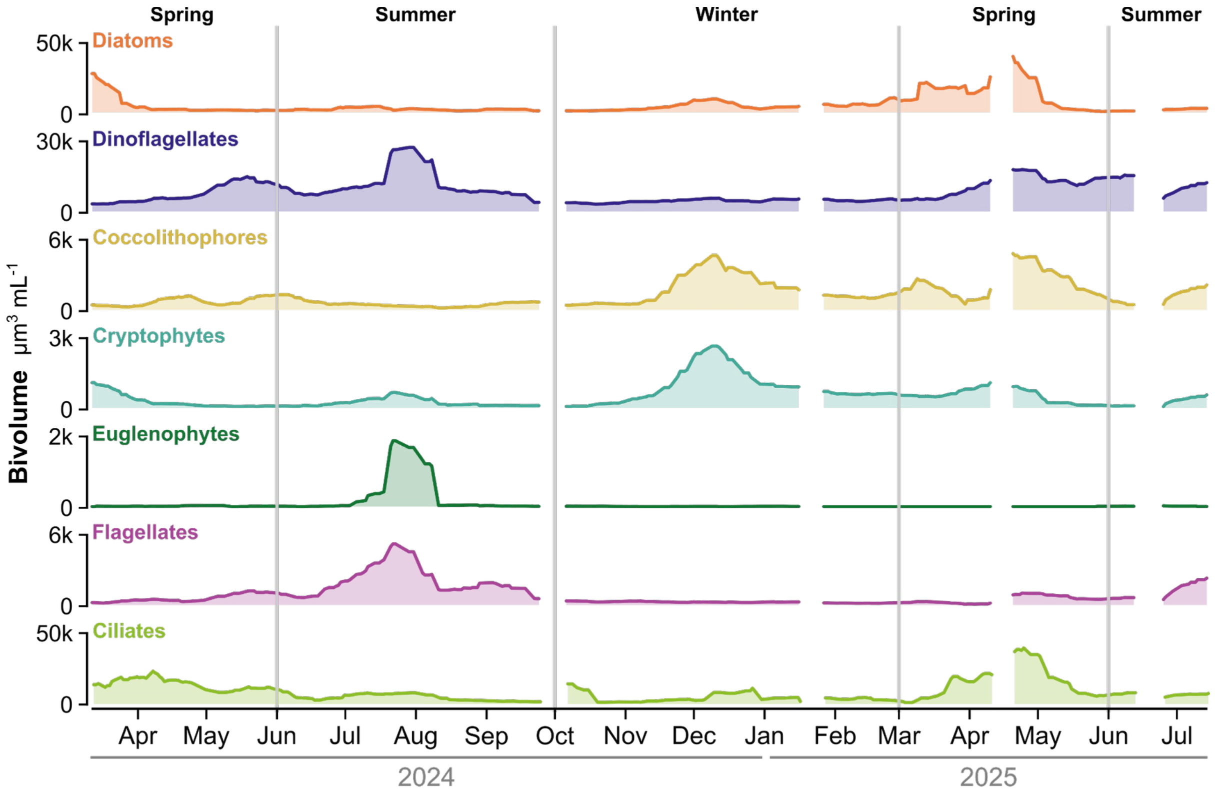


**Figure S3.** Smoothed estimated biovolume of major protist groups in the Gulf of Aqaba. Lines are centered 21-day rolling means calculated from observations within 10 days before and after each date. Sampling gaps longer than 6 days are left unconnected. Note the difference in y-axis between groups.

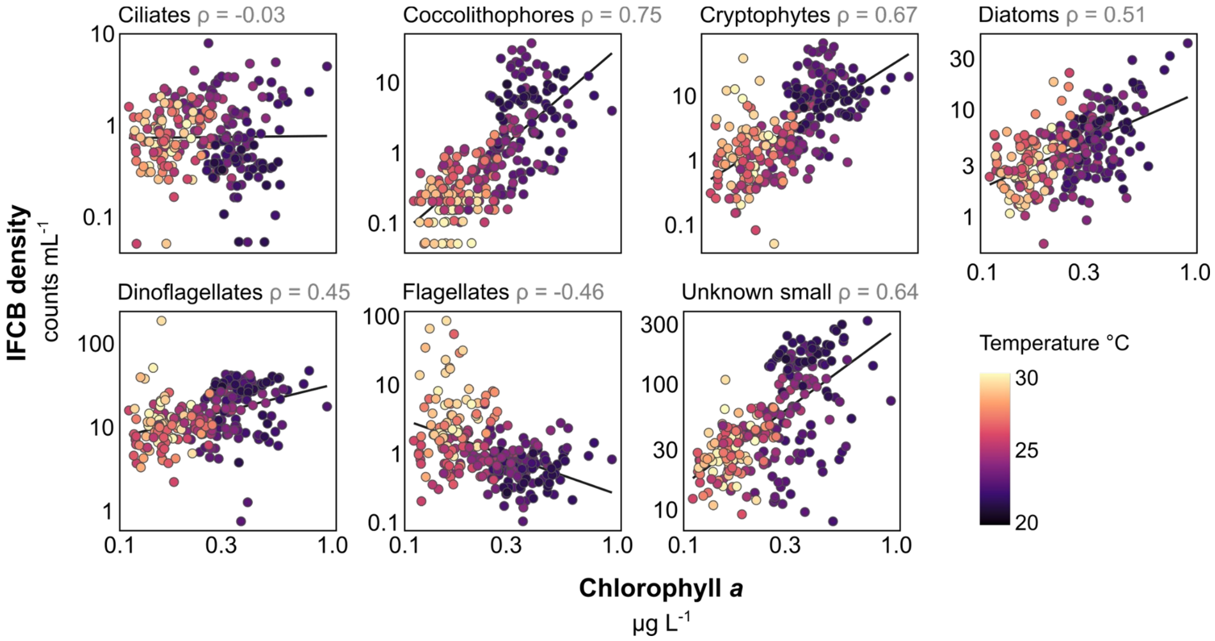


**Fig. 5.** Comparisons between the concentration of chlorophyll *a* and IFCB-derived cell density for the seven dominant protist groups. Chlorophyll *a* was measured daily at station 2 and IFCB data are from station 1 (n = 271). The stations are separated by 400 m (see Figure 1A). Both axes are $\log_{10}$-scaled. Point color indicates sea-surface temperature (°C). Each panel reports Spearman's rho between chlorophyll *a* and density of the group. Correlations were positive and significant for Coccolithophores, Cryptophytes, Diatoms, Dinoflagellates, and Unknown small ($p < 0.001$), negative for Flagellates ($p < 0.001$), and absent for Ciliates. Note the difference in y-axis between panels.
